# The leucine-rich repeat (LRR) domain of NLRP3 is required for NLRP3 inflammasome activation in macrophages

**DOI:** 10.1101/2022.05.25.493460

**Authors:** Yanhui Duan, Jihong Wang, Juan Cai, Nathan Kelley, Yuan He

**Author notes:** Address correspondence to Dr. Yuan He, Department of Biochemistry, Microbiology, and Immunology, Wayne State University School of Medicine, 540 E Canfield Ave, Detroit, MI 48201. Yanhui Duan: School of Pharmaceutical Science and Technology, Hangzhou Institute for Advanced Study, University of Chinese Academy of Sciences, Hangzhou, China, 310024.

## Abstract

The NLRP3 inflammasome is a critical component of innate immunity that defends the host from microbial infections. However, its aberrant activation contributes to the pathogenesis of several inflammatory diseases. Activation of the NLRP3 inflammasome induces the secretion of proinflammatory cytokines IL-1β and IL-18, and pyroptotic cell death. NLRP3 contains a leucine-rich repeat (LRR) domain at its C-terminus. Although posttranslational modifications in this LRR domain have been shown to regulate NLRP3 inflammasome activation, the role of the entire LRR domain in NLRP3 inflammasome activation remains controversial. Here, we generated mouse macrophages that express an endogenous NLRP3 mutant lacking the LRR domain. Deletion of the LRR domain destabilized endogenous NLRP3 protein and abolished NLRP3 inflammasome activation in macrophages. Furthermore, using NLRP3-deficient macrophages that are reconstituted with NLRP3 mutants lacking the LRR domain, we found that deletion of the LRR domain inhibited NLRP3 inflammasome activation. Mechanistically, deletion of the LRR domain abolished NLRP3 self-association, oligomerization, and interaction with the essential regulator NEK7. Our results demonstrate a critical role for the LRR domain in NLRP3 inflammasome activation.

## Introduction

Inflammasomes are cytoplasmic multi-protein complexes that form in response to microbial infection or cell damage (1). In most cases, an inflammasome contains a pattern recognition receptor as the sensor, the adaptor protein apoptosis-associated speck-like protein containing a caspase-activation and recruitment domain (ASC), and the inflammatory caspase-1 (2,3). Inflammasome assembly results in caspase-1 autocleavage and activation. Activated caspase-1 processes the proinflammatory cytokines pro-interleukin (IL)-1β and pro-IL-18 into their mature forms, and cleaves gasdermin D (GSDMD) to release its active N-terminus, which forms pores in the plasma membrane to induce cytokine release and pyroptotic cell death (4-6). To date, six inflammasomes have been well documented, including the nucleotide-binding oligomerization domain (NOD)-like receptor (NLR) family members NLRP1, NLRP3, NLRP6, NLRC4, pyrin, and absent in melanoma 2 (AIM2) inflammasomes (2,7). Among these inflammasomes, the NLRP3 inflammasome has been intensively investigated due to its protective role in host defense against various infections as well as its pathogenic role in several inflammatory disorders (8,9).

The mechanism underlying NLRP3 inflammasome activation remains not fully understood. A diverse spectrum of stimuli, including ATP, pore-forming toxin (e.g., nigericin), and particulate matter (e.g., monosodium urate crystals, amyloids, and silica) have been shown to activate the NLRP3 inflammasome (10-12). Therefore, it is unlikely that NLRP3 directly recognizes these stimuli. Instead, molecular or cellular signal events downstream of these stimuli, such as altered intracellular ion homeostasis, mitochondrial dysfunction, reactive oxygen species (ROS) production, and lysosomal damage, have been proposed to trigger NLRP3 inflammasome activation (10-12). Most NLRP3 stimuli trigger potassium efflux, which has been considered a critical signal for NLRP3 inflammasome activation (13,14). However, how NLRP3 senses the drop in intracellular potassium concentration remains unknown. Furthermore, several NLRP3-interacting proteins including HSP90, SGT1, TXNIP, NEK7, DDX3X, and RACK1, have been shown to regulate NLRP3 inflammasome activation (15-20). However, the molecular mechanism of these regulators remains to be further defined.

NLRP3 consists of three domains: an N-terminal pyrin domain (PYD), a central NAIP, CIITA, HET-E, and TP1 (NATCH) domain, and a C-terminal LRR domain (21). NLRP3 interacts with the adaptor protein ASC through PYD-PYD interactions to induce the formation of filamentous aggregates known as ASC specks (22,23). The NATCH domain has an adenosine triphosphatase activity required for NLRP3 conformational change and oligomerization (24,25). In contrast, the role of the LRR domain in NLRP3 inflammasome activation is more complex. Earlier studies have suggested that the LRR domain has an autoinhibitory function through intramolecular interactions (26). Recently, several posttranslational modifications, including phosphorylation and ubiquitination, have been found in the LRR domain to regulate NLRP3 inflammasome activation (27-30). However, there are conflicting reports on whether the entire LRR domain plays a role in NLRP3 inflammasome activation. Some studies have shown that the replacement of the LRR domain by the lacZ protein or its deletion abolished NLRP3 inflammasome activation (31,32). Furthermore, structural studies have shown that the LRR domain mediated the NLRP3-NEK7 interaction and the formation of the NLRP3 cage structure, which disperses the *trans*-Golgi network at the early stage of the inflammasome pathway (33,34). However, others have shown that the LRR domain is dispensable for NLRP3 inflammasome activation (35,36).

In this study, we have used mouse knock-in macrophages that express an endogenous NLRP3 mutant lacking the LRR domain, and NLRP3-deficient macrophages that are reconstituted with NLRP3 mutants lacking the LRR domain. Our results demonstrate that the LRR domain promotes the stability of endogenous NLRP3 and is required for NLRP3 inflammasome activation. Furthermore, our results show that the LRR domain mediates NLRP3 self-association, oligomerization, and interaction with its essential regulator NEK7.

## Results

### Deletion of the LRR domain destabilizes endogenous NLRP3 protein and abrogates nigericin-induced NLRP3 inflammasome activation in macrophages

Previous studies showed conflicting results on the role of the LRR domain in NLRP3 inflammasome activation. Notably, most of these studies overexpressed NLRP3 mutants lacking the LRR domain by transfection or viral infection, or replaced the LRR domain of the endogenous NLRP3 protein with a large LacZ protein, which may interfere with NLRP3 function (31,32,35,36). To determine the exact role of this LRR domain in NLRP3 inflammasome activation, we used the CRISPR-Cas9 gene-editing system to generate immortalized mouse bone marrow-derived macrophages (iBMDMs) that endogenously express an NLRP3 mutant lacking the LRR domain. An NLRP3 mutant [NLRP3(1-720)] was chosen as it lacks the entire LRR domain and has been reported to fully restore NLRP3 inflammasome activation in NLRP3-deficient mouse macrophages (Fig. 1*A*) (35). We selected three knock-in (KI) macrophage clones (*Nlrp3*^*C721Stop/C721Stop*^) in which a stop codon (TGA) has successfully replaced the cysteine (C721) codon (TGC) at the coding region of mouse *Nlrp3* gene (Fig. 1*B*). These KI macrophages are expected to express an NLRP3 mutant that consists of the first 720 amino acid residues and lacks the entire LRR domain.

**Figure 1.**
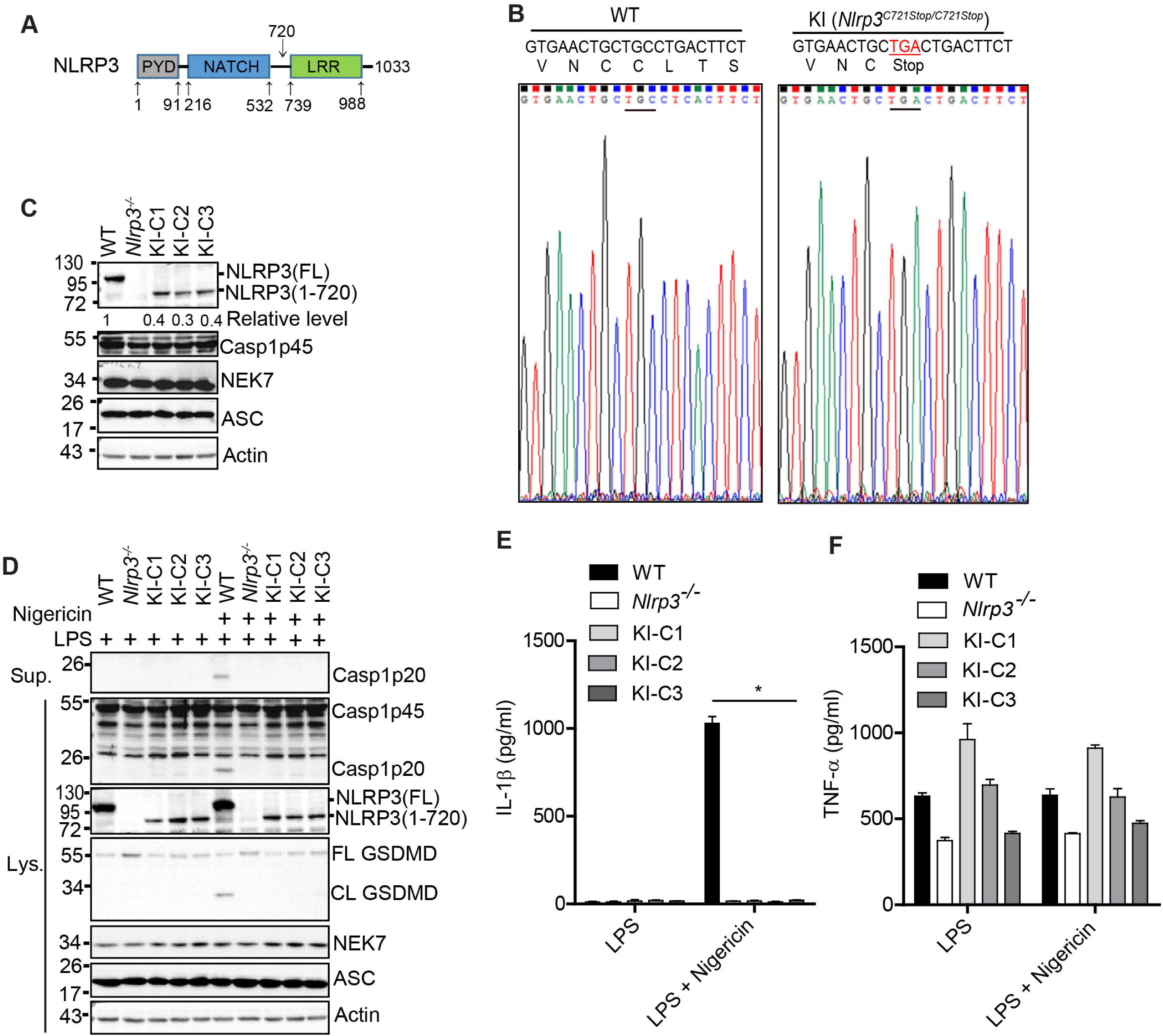
The LRR domain promotes the stability of endogenous NLRP3 and is required for nigericin-induced NLRP3 inflammasome activation in macrophages. *A*, Schematic presentation for domains of mouse NLRP3 (UniProtKB: Q8R4B8). The bottom numbers show the positions of amino acid residues at the start and end of each domain. The top number (720) shows the position of the last amino acid residue for an NLRP3 LRR deletion mutant [NLRP3 (1-720)]. *B*, sequencing verification for the replacement of a cysteine codon (TGC) by a stop codon (TGA) in mouse *Nlrp3* gene. *C*, immunoblot analysis of NLRP3 inflammasome components in wild-type (WT), *Nlrp3*^*-/-*^, and knock-in macrophages (three individual clones: KI-C1, KI-C2, KI-C3). Actin was used as a control. Representative blots (n = 2). The band density of NLRP3 was measured by ImageJ and normalized to the full-length NLRP3. *D*, macrophages were stimulated with LPS (4 h) alone or plus 5 μM nigericin (1 h). Cell lysates and supernatants were immunoblotted with indicated antibodies. Representative blots (n = 3). Measurement of IL1-β (*E*) and TNF-α (*F*) in the supernatants from stimulated macrophages by ELISA. Representative data(n=3). \**p* < 0.05 (one-way ANOVA). Data are the mean ± SEM of triplicate wells. FL, full-length. CL, cleaved.

We assessed the protein expression of endogenous NLRP3 protein from both wild-type (parent cells) and KI macrophages by western blot. An antibody that recognized an epitope in the N-terminal pyrin domain of NLRP3 protein (Cryo-2) was used to detect both full-length and truncation mutant NLRP3 from these macrophages. In addition, macrophages derived from *Nlrp3*^*-/-*^ mice were included as controls in our experiments. The full-length NLRP3 (∼118 kDa) was ready detected in wild-type macrophages at the resting state (Fig. 1*C*). As expected, all three KI macrophage clones expressed the short version of NLRP3 protein (∼ 79 kDa), further confirming our sequencing results (Fig. 1*C*). Of note, no protein band was shown up in the lane for *Nlrp3*^*-/-*^ macrophages, indicating the specificity of our antibody (Fig.1*C*). Importantly, the detected protein level of this NLRP3 mutant from each clone was markedly lower (less than 50%) when compared with that of full-length NLRP3 expressed in wild-type macrophages (Fig.1*C*). However, the expression levels of other NLRP3 inflammasome components, including caspase-1, NEK7, and ASC, were comparable between wild-type and KI macrophages (Fig.1*C*). Next, we assessed NLRP3 inflammasome activation in these macrophages under the conditions of LPS priming or LPS priming plus nigericin treatment. Previous studies have suggested that the LRR domain keeps an inactive conformation of NLRP3 through intramolecular interaction (26). Similar to wild-type macrophages, KI macrophages did not exhibit caspase-1 activation, GSDMD cleavage, or IL-1β secretion after LPS priming (Fig.1, *D* and *E*). Nigericin, a classic NLRP3 stimulus, induced NLRP3 inflammasome activation in wild-type macrophages, as revealed by caspase-1 activation, GSDMD cleavage, and IL-1β secretion (Fig.1, *D* and *E*). In contrast, KI macrophages, similar to NLRP3-deficient macrophages, were defective in NLRP3 inflammasome activation after LPS priming plus nigericin treatment (Fig.1, *D* and *E*). The secretion of TNF-α, an inflammasome-independent cytokine, was comparable among these macrophages (Fig.1*F*). Collectively, these results indicate that the LRR domain is required for endogenous NLRP3 stability and nigericin-induced NLRP3 inflammasome activation in macrophages.

### Deletion of the LRR domain inhibits NLRP3 inflammasome activation by diverse NLRP3 stimuli

NLRP3 inflammasome activation is induced by potassium efflux-dependent (e.g, ATP, nigericin, Nano-SiO_2_) or independent stimuli (e.g., Imiquimod) (11,12). Given our data showing that KI macrophages were defective in nigericin-induced NLRP3 inflammasome activation, we next examined whether this defect was common to other NLRP3 stimuli. As they secreted a similar level of TNF-α after LPS priming, the KI macrophage clone KI-C2 (KI henceforth) and wild-type macrophages were used for the following experiments (Fig.1*F*). Macrophages were stimulated with ATP, nigericin, Nano-SiO_2_, or Imiquimod after LPS priming. KI macrophages showed markedly reduced caspase-1 activation, GSDMD cleavage, and IL-1β secretion when compared to wild-type macrophages (Fig.2, *A* and *B*). In contrast, KI macrophages had comparable AIM2 inflammasome activation induced by poly(dA:dT) or NLRC4 inflammasome activation by *Salmonella* when compared to wild-type macrophages (Fig.2, *A* and *B*). Inflammasome activation induces the formation of ASC aggregates called ASC specks, which are composed of ASC oligomers in the cytosol (22). Both ATP and nigericin induced ASC speck formation in wild-type macrophages, whereas almost no ASC speck was detected in KI macrophages for both stimuli (Fig.2, *C* and *D*). However, KI macrophages had comparable ASC speck formation in response to *Salmonella* as compared to wild-type macrophages (Fig.2, *C* and *D*). Biochemical analysis revealed that KI macrophages had much less ASC oligomerization as compared to wild-type cells after treatment of ATP or nigericin, although they showed a comparable level of ASC oligomerization in response to *Salmonella* (Fig.2*E*). To exclude whether other nonspecific factors besides this LRR domain deletion in NLRP3 attributed to the defect of NLRP3 inflammasome activation observed in KI macrophages, we expressed C-terminal HA-tagged NLRP3 full-length [NLRP3(FL)], LRR deletion mutant [NLRP3(1-720)], or the LRR domain [NLRP3(730-1033)] in KI macrophages by a lentiviral vector and assessed NLRP3 inflammasome activation in response to nigericin. Expression of full-length NLRP3 restored NLRP3 inflammasome activation, indicated by caspase-1 activation and IL-1β secretion, in KI macrophages, while both the LRR deletion mutant and LRR domain alone failed to rescue NLRP3 inflammasome activation (Fig.2, *F* and *G*). Taken together, these results indicated that the LRR domain is required for NLRP3 inflammasome activation in macrophages.

**Figure 2.**
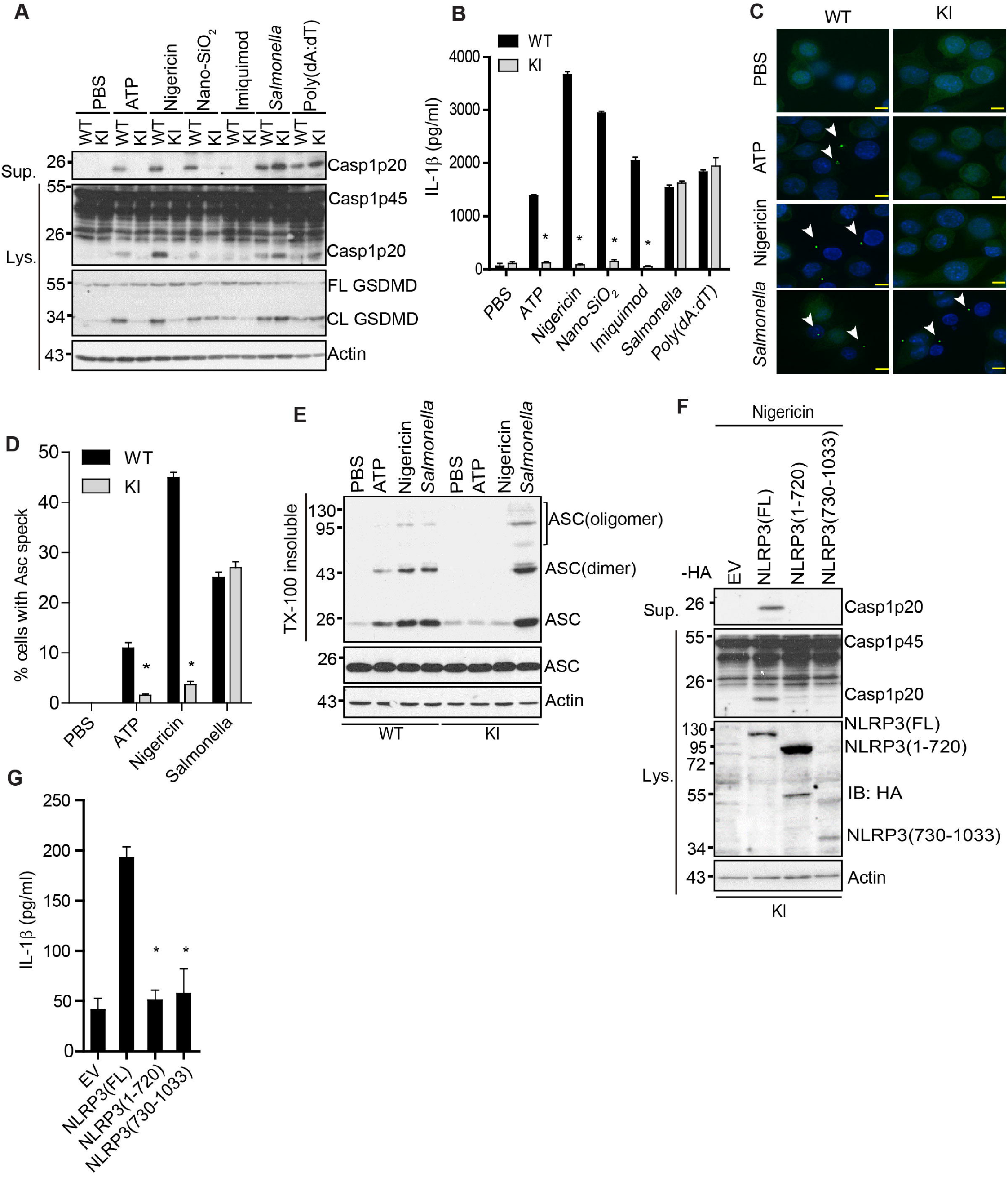
The LRR domain is required for NLRP3 inflammasome activation induced by diverse stimuli. *A*, LPS-primed WT and KI macrophages were primed with LPS (4 h) and then stimulated with PBS (mock), 5 mM ATP (1 h), 5 μM nigericin (1 h), 200 μg/ml Nano-SiO_2_ (4 h), 15 μg/ml Imiquimod (4 h), 4 μg/ml poly(dA:dT) (4 h) or *Salmonella* (m.o.i= 10, 2h). Cell lysates and supernatants were immunoblotted with indicated antibodies. Representative blots (n = 3). *B*, Measurement of IL-β in the supernatants from (A) by ELISA. Representative data (n=3). \**p* < 0.05 (unpaired two-sided t-test). Data are the mean ± SEM of triplicate wells. *C*, ASC immunostaining in wild-type or KI macrophages primed with LPS and stimulated with PBS, ATP, nigericin, or *Salmonella*. Scale bars, 10 μm. ASC specks are indicated by white arrows. Representative images (n=3). *D*, Quantification of ASC specks from (C). The percentage of ASC speck-containing cells was calculated from three different fields with at least 100 cells each. \**p* < 0.05 (unpaired two-sided t-test). *E*, immunoblot analysis of ASC oligomerization from Triton X (TX)-100 insoluble and soluble fractions in WT and KI macrophages. *F*, KI macrophages were transduced with a lentiviral vector expressing c-terminal HA-tagged full-length (FL), LRR domain deletion mutant [NLRP3 (1-720)], or the LRR domain alone [NLRP3 (730-1033)]. Macrophages were primed with LPS and then stimulated with nigericin. Cell lysates and supernatants were immunoblotted with indicated antibodies. Actin was used as a control. Representative blots (n=3). *G*, Measurement of IL-β in the supernatants from (*F*) by ELISA. \**p* < 0.05 (unpaired two-sided t-test). Data are the mean ± SEM of triplicate wells. FL, full-length. CL, cleaved.

### NLRP3 mutants lacking the LRR domain fail to restore NLRP3 inflammasome activation in NLRP3-deficient macrophages

Several LRR domain-lacking mutants of NLRP3, including NLRP3 1-686 and 1-720, have been reported to fully restored inflammasome activation in mouse NLRP3-deficient macrophages (35). As shown in our data, endogenously expressed protein NLRP3(1-720) was less stable and its level was lower than the full-length NLRP3 in macrophages. As a complementary approach, we constitutively expressed full-length NLRP3, NLRP3 mutant 1-686, or 1-720 in mouse NLRP3-deficient macrophages and selected macrophage clones that express comparable protein levels of NLRP3 (Fig.3*A*). In contrast to the previous study in which macrophages were primed with LPS for 11 hours in the presence of doxycycline, we primed reconstituted macrophages with LPS for 4 hours before stimulation with ATP, nigericin, Nano-SiO_2_, or Imiquimod (35). Consistent with our results from KI macrophages, macrophages reconstituted with NLRP3 mutant 1-686 or 1-720 showed a significant reduction in caspase-1 activation, GSDMD cleavage, and IL-1β secretion when compared to macrophages with full-length NLRP3 (Fig.3, *A* and *B*). The levels of TNF-α secretion were comparable between macrophages reconstituted with wild-type or mutant NLRP3 under the same conditions (Fig.3*C*). Accordingly, macrophages reconstituted with NLRP3 mutant 1-686 or 1-720 had less ASC oligomerization in response to ATP or nigericin when compared to macrophages with full-length NLRP3 (Fig.3*D*). These results further indicated that the LRR domain is also essential for NLRP3 inflammasome activation in reconstituted NLRP3-deficient macrophages.

**Figure 3.**
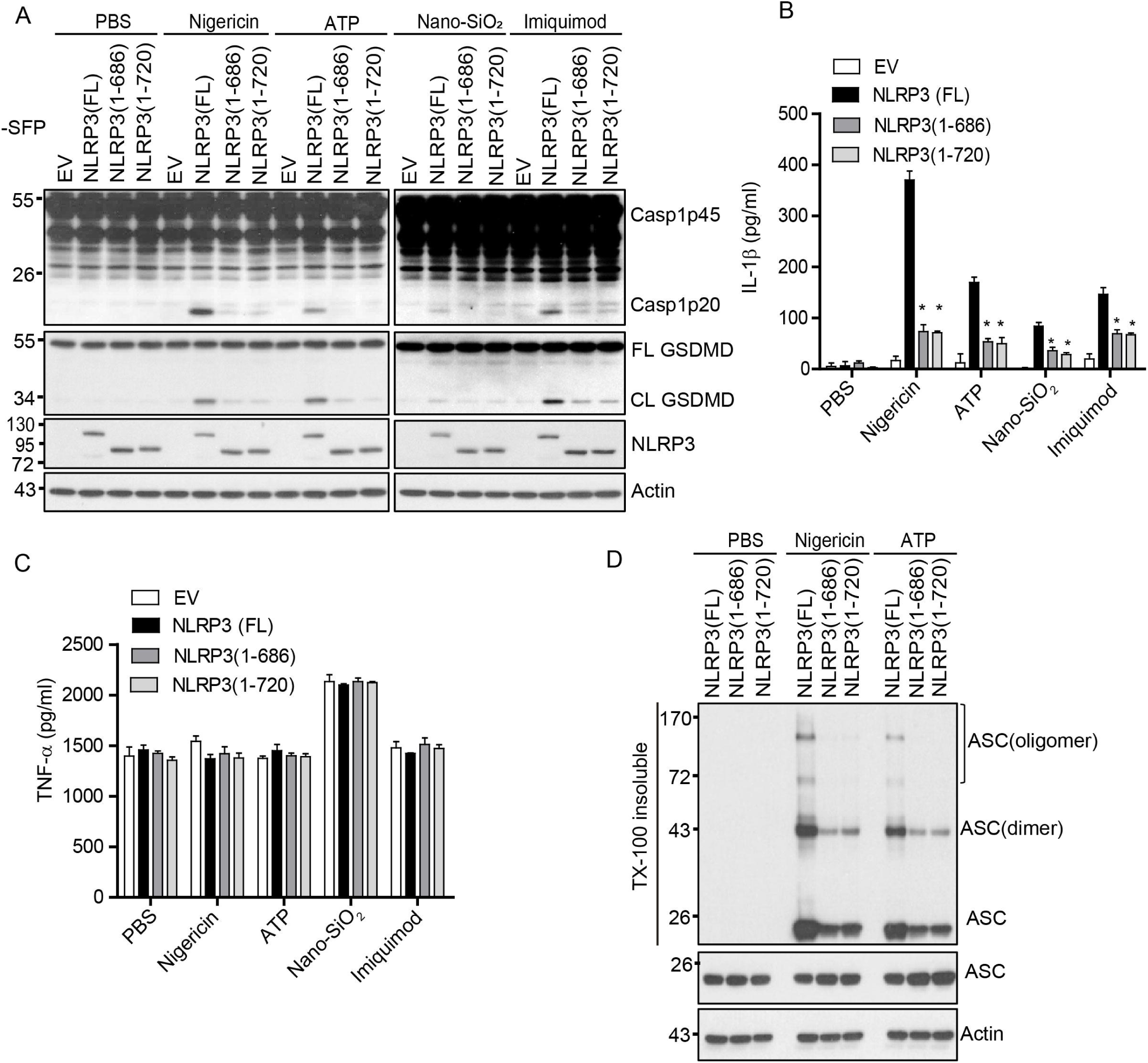
LRR domain deletion mutants of NLRP3 fail to fully rescue NLRP3 inflammasome activation in mouse *Nlrp3*^*-/-*^ macrophages. A, mouse *Nlrp3*^*-/-*^ macrophages were transduced with a lentiviral vector expressing c-terminal SFP-tagged full-length (FL), LRR domain deletion mutant 1-686 [NLRP3 (1-686)], or LRR domain deletion mutant 1-720 [NLRP3 (1-720)]. LPS-primed macrophages were stimulated with PBS (mock), 5 μM nigericin (1 h), 5 mM ATP (1 h), 200 μg/ml Nano-SiO_2_ (4 h), or 15 μg/ml Imiquimod (4 h). Mixtures of cell lysates and supernatants were immunoblotted with indicated antibodies. Measurement of IL-β (B) and TNF-α (C) in the supernatants from stimulated macrophages by ELISA. D, immunoblot analysis of ASC oligomerization from Triton X (TX)-100 insoluble and soluble fractions in macrophages expressing indicated full-length or LRR deletion mutant of NLRP3. Representative blots (n=3). Data are the mean ± SEM of triplicate wells. \**p* < 0.05 (unpaired two-sided t-test). EV, empty vector. FL, full-length. CL, cleaved.

### Deletion of the LRR domain inhibits NLRP3 self-association and oligomerization

Recent structural studies with purified NLRP3 oligomers have revealed that NLRP3 self-associates through the LRR-LRR interaction (34,37). To investigate how the LRR domain controls NLRP3 inflammasome activation, we assessed the effects of the LRR domain deletion on NLRP3 self-association and oligomerization in cells. We co-expressed SFP or HA-tagged full-length or NLRP3 mutants in HEK 293T cells. The full-length NLRP3 showed self-association as SFP-tagged NLRP3 pulled down HA-tagged NLRP3. In contrast, self-association for both NLRP3 mutants 1-686 and 1-720 was markedly reduced (Fig.4*A*). In macrophages, after stimulation of NLRP3 stimuli, NLRP3 forms high-molecular-mass complexes, as indicated by protein native gels (17). We compared these high-molecular-mass forms of NLRP3 in cells reconstituted with full-length or the LRR deletion mutant. Macrophages reconstituted with the full-length NLRP3 displayed the high-molecular-mass NLRP3 complexes (>1200 kDa) after stimulation of ATP and nigericin. In contrast, these high-molecular-mass forms of NLRP3 were markedly reduced in macrophages reconstituted with NLRP3 truncation mutant 1-686 or 1-720 (Fig.4*B*). Accordingly, in 2-D native gels, these high-molecular-mass forms of NLRP3 proteins were markedly reduced in macrophages reconstituted with NLRP3 mutant 1-686 or 1-720 as compared with the full-length NLRP3 (Fig.4*C*). Taken together, our results indicate that the LRR domain mediates NLRP3 self-association and oligomerization.

**Figure 4.**
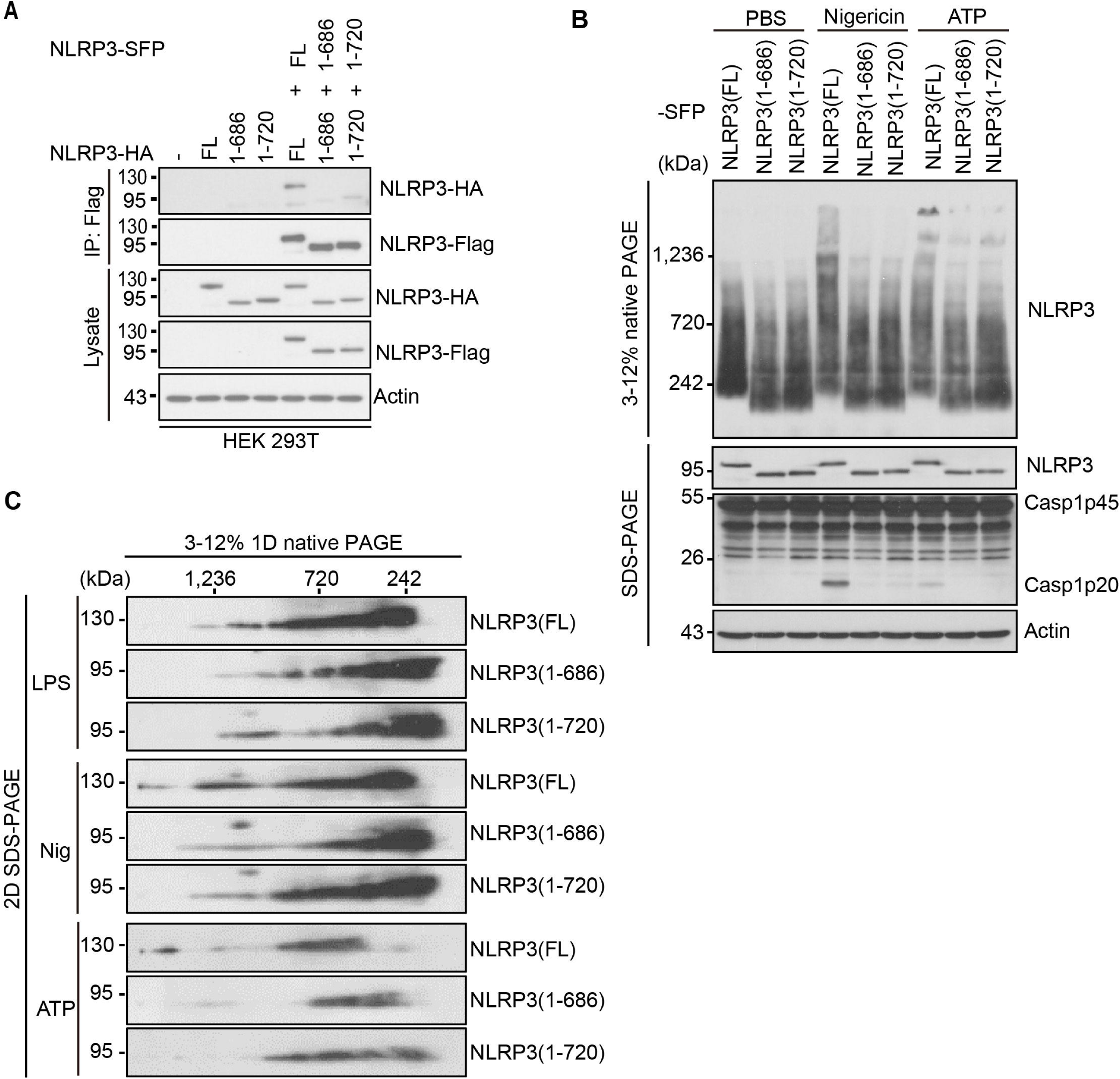
LRR domain deletion mutants of NLRP3 show defects in NLRP3 self-association and oligomerization. *A*, HA-tagged mouse full-length NLRP3, LRR domain deletion mutant NLRP3 (1-686), or NLRP3 (1-720)] was expressed alone or with indicated SFP-tagged protein in HEK293T cells. Cell lysates were immunoprecipitated with anti-Flag antibody and immunoblotted with indicated antibodies. *B*, mouse *Nlrp3*^*-/-*^ macrophages were reconstituted with SFP-tagged wild-type or mutant NLRP3 and stimulated with PBS (mock), ATP, or nigericin after LPS priming. NLRP3 oligomerization was analyzed by blue native PAGE and immunoblotting with an anti-Flag antibody. Cell lysates were also analyzed by SDS–PAGE and immunoblotting with indicated antibodies. *C*, macrophage cell lysates were separated by the blue native PAGE, followed by a second dimension of SDS–PAGE and western blot. Representative blots (n=3). FL, full-length.

### Deletion of the LRR domain abolishes the NEK7-NLRP3 interaction

Previous studies have shown that the LRR domain in NLRP3 is required for the NEK7-NLRP3 interaction in HEK 293T cells (17,18). Furthermore, a structural study of the NEK7-NLRP3 complex has shown that NEK7 interacts with the LRR domain of NLRP3 (33). Surprisingly, NLRP3 mutants lacking the LRR domain including 1-686 were reported to pull down NEK7 in macrophages (35). Therefore, we examined whether the LRR domain is required for NLRP3-NEK7 interaction in our reconstituted macrophages. Both NLRP3 mutants 1-686 and 1-720 failed to pull down NEK7 from macrophages stimulated LPS or LPS plus ATP (Fig.5*A*). The NLRP3 mutant 1-720 or the LRR domain alone also failed to pull down NEK7 from HEK 293T cells (Fig.5*B*). According to the Uniprot/Swiss-Prot, there are 9 leucine-rich repeats in the LRR domain of NLRP3. To further define the exact leucine-rich repeat(s) of the LRR domain that is essential for NLRP3-NEK7 interaction, we generated a series of NLRP3 mutants which progressively lack one leucine-rich repeat starting from the C-terminus. Co-expression of the Flag-tagged NLRP3 mutant and HA-tagged NEK7 in HEK 293T cells was used to assess the interaction between NEK7 and NLRP3 mutants. All these NLRP3 mutants that lack one or several leucine repeats in the LRR domain lost the ability for NEK7 binding (Fig.5*C*). Collectively, these results indicate that the intact LRR domain is essential for NEK7-NLRP3 interaction.

**Figure 5.**
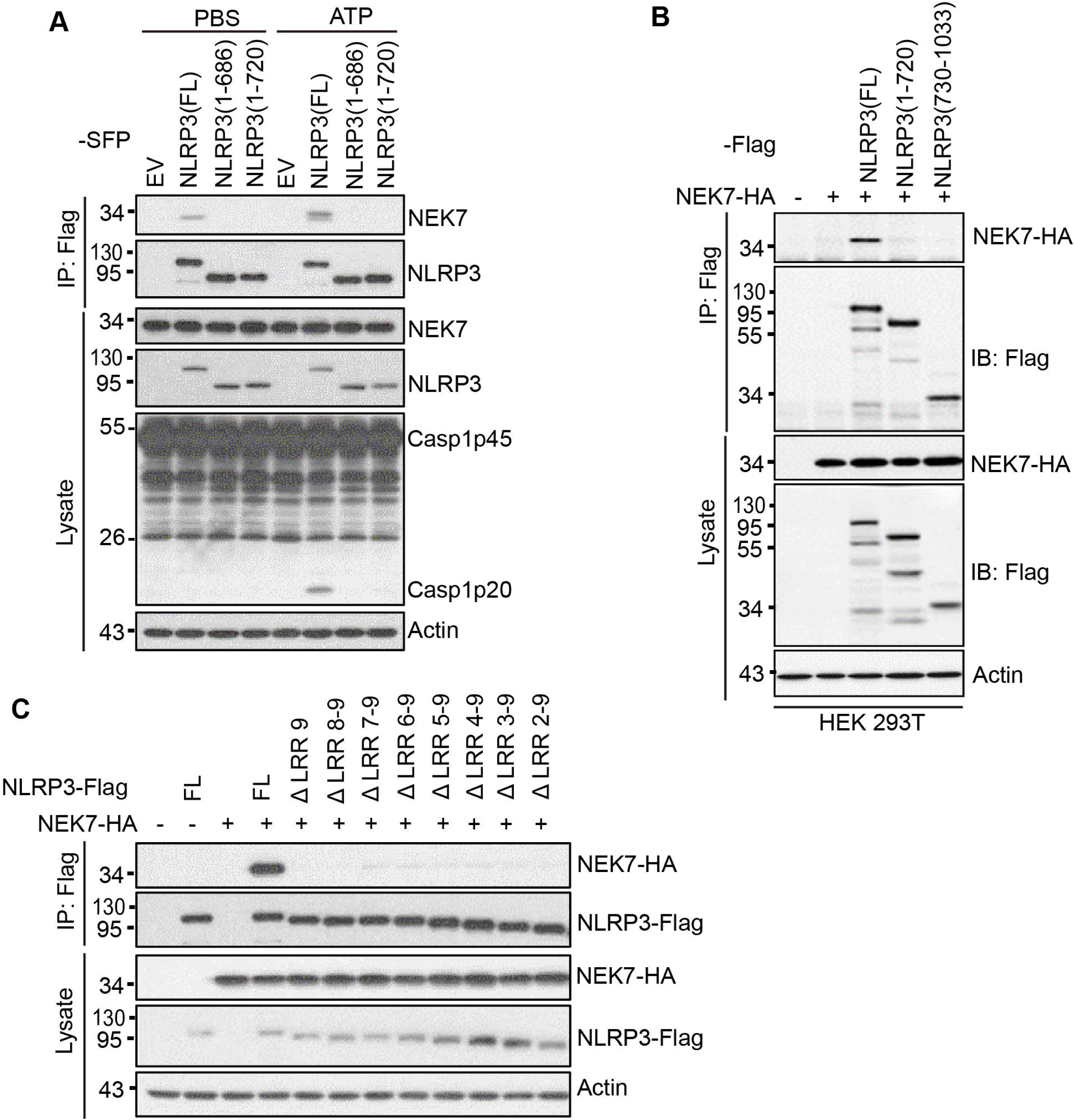
Deletion of the LRR domain in NLRP3 abrogates NEK7-NLRP3 interaction. A, mouse *Nlrp3*^*-/-*^ macrophages were reconstituted with SFP-tagged full-length or mutant NLRP3 and stimulated with PBS (mock) or 5 mM ATP (1 h) after LPS priming. Cell lysates were immunoprecipitated with anti-Flag antibody and immunoblotted with indicated antibodies. B, Flag-tagged full-length, NLRP3 (1-720), or NLRP3 (730-1033) was co-expressed with HA-tagged NEK7 in HEK 293T cells. Cell lysates were immunoprecipitated with anti-Flag antibody and immunoblotted with indicated antibodies. C, Flag-tagged NLRP3 mutants with deletion of one or multiple leucine-repeats were co-expressed with HA-tagged NEK7 in HEK 293T cells. Cell lysates were immunoprecipitated with anti-Flag antibody and immunoblotted with indicated antibodies. ΔLRR 9, NLRP3 (1-964); ΔLRR 8-9, NLRP3 (1-936); ΔLRR 7-9, NLRP3 (1-907); ΔLRR 6-9, NLRP3 (1-879); ΔLRR 5-9, NLRP3 (1-850); ΔLRR 4-9, NLRP3 (1-818); ΔLRR 3-9, NLRP3 (1-793); ΔLRR 2-9, NLRP3 (1-765). Representative blots (n=3). EV, empty vector. FL, full-length.

## Discussion

NLRP3 activation triggers the assembly of an inflammasome that plays a critical role in innate immunity. In addition to its N-terminal pyrin domain and central NATCH domain, NLRP3 contains an LRR domain at its C-terminus. Several post-translational modifications have been found on the LRR domain to regulate NLRP3 inflammasome activation (11,12). In this study, we investigated the role of this LRR domain in NLRP3 inflammasome activation in mouse macrophages. Our results show that the LRR domain promotes the stability of endogenous NLRP3 protein and is required for NLRP3 inflammasome activation in macrophages. Furthermore, we show that the LRR domain mediates NLRP3 self-association, oligomerization, and interaction with NEK7, which are critical molecular events for NLRP3 inflammasome activation.

The stability of a protein is essential for its function in cells. To our best knowledge, the role of the LRR domain in endogenous NLRP3 stability has not been addressed yet as most of the NLRP3 LRR deletion mutants were overexpressed by a viral infection or plasmid transfection in previous studies. In contrast, we used knock-in macrophages that express an endogenous NLRP3 mutant that lacks the entire LRR domain. Our results suggest that the LRR domain is required for NLRP3 stability in macrophages. This role of the LRR domain in NLRP3 protein stability, although it has not been explored further, has been suggested in previous studies (31). In addition, previous studies report that the chaperon heat-shock protein 90 (HSP90) interacts with NLRP3 via the LRR domain and stabilizes the NLRP3 protein (15). Therefore, we hypothesize that the deletion of the LRR domain in NLRP3 might cause the loss of this HSP90-mediated protection and thus render its degradation by the proteasome. In support of this hypothesis, the E3 ubiquitin ligase Cbl-b has been reported to ubiquitinate the NATCH domain at K496 for proteasome-mediated degradation of NLRP3 (28). Interestingly, short NLRP3 transcripts that encode NLRP3 truncation forms have been reported in mouse and human cells (38,39). It remains unknown whether those truncation proteins of NLRP3 are unstable in cells. In addition, most of the Nod-like receptors contain an LRR domain at their C-terminus (40). It remains to be investigated whether the LRR domain has a similar role in their stability.

There are conflicting reports on the role of the LRR domain in NLRP3 inflammasome activation. Previous studies have suggested that the LRR domain plays an inhibitory role in NLRP3 inflammasome activation (26). However, we did not observe constitutive activation of the NLRP3 inflammasome in mouse macrophages expressing an endogenous or exogenous LRR domain-lacking NLRP3 mutant. Recently, Hafner-Bratkovič et al. reported that mouse NLRP3 truncation mutants including 1-686 and 1-720 fully restored NLRP3 inflammasome activation in NLRP3-deficient macrophages, suggesting a dispensable role for this LRR domain in NLRP3 inflammasome activation (35). In contrast, Niu et al. reported that an NLRP3 mutant 1-688 (mouse 1-686) failed to restore NLRP3 inflammasome activation in NLRP3-deficient human monocytic U937 cells (32). In addition, BMDMs expressing an NLRP3 mutant, in which the LRR domain was replaced by a LacZ protein, were defective in NLRP3 inflammasome activation induced by monosodium urate crystals (31). Consistent with the latter two studies, our results indicate that the LRR domain is required for NLRP3 inflammasome activation in mouse macrophages. In our knock-in macrophages that express an endogenous NLRP3 mutant lacking the LRR domain, NLRP3 inflammasome activation was completely abrogated in response to all tested NLRP3 stimuli. This defect may also partially be attributed to the low protein level of this endogenous NLRP3 mutant in knock-in macrophages. However, overexpression of this NLRP3 mutant failed to restore NLRP3 inflammasome activation in these knock-in macrophages, suggesting an intrinsic role for the LRR domain in NLRP3 inflammasome activation. In addition, macrophages expressing NLRP3 truncation mutant 1-686 or 1-720, to a similar level as the full-length protein, were defective in NLRP3 inflammasome activation in our reconstitution experiments. We do not know the exact reasons for the above disparities. One possibility is that Hafner-Bratkovič et al. induced a high level of NLRP3 proteins by using a longer time (11 hours) of LPS priming in a doxycycline-inducible system (35). This experimental setting might induce a high level of protein expression and thus bypass the requirement of the LRR domain in NLRP3 inflammasome activation. This is likely also applied to the reconstituted NLRP3 inflammasome in HEK293T cells (36). In contrast, we selected reconstituted NLRP3-deficient macrophage clones that expressed a similar level of NLRP3 protein as the wild-type macrophages, and primed cells for a short time (4 hours) before adding NLRP3 stimuli. Importantly, we found that LRR deletion reduced NLRP3 self-association and oligomerization. Consistent with this observation, NLRP3 mutants with mutations in the LRR domain have been reported to be defective in the formation of a double-ring cage structure of mouse NLRP3 (34). Therefore, it is conceivable that deletion of the LRR will abolish the NLRP3 cage formation and subsequent inflammasome activation. Additionally, we further confirm and extend previous findings that an intact LRR domain in NLRP3 is required for its interaction with NEK7 (17,18). Since the NLRP3 cage structure is not compatible with NEK7 binding, it remains to be determined when and where the LRR domain is involved in the interaction with NEK7 in this inflammasome activation pathway.

In summary, our findings demonstrated a critical role for the LRR domain in endogenous NLRP3 protein stability and inflammasome activation in macrophages. Our results reveal that the LRR domain controls inflammasome activation by mediating NLRP3 self-association, oligomerization, and interaction with NEK7.

## Experimental procedures

### Antibodies and Reagents

Anti-Flag (A00187-200), Anti-HA (A01244-100), Anti-Actin (A00730-100), and protein G resin (L00209) were purchased from GenScript. Anti-NLRP3 (AG-20B-0014-C100) was purchased from Adipogen. Anti-GSDMD (ab209845) and Anti-NEK7 (ab133514) were purchased from Abcam. Anti-ASC (67824) was purchased from CST. Anti-mouse Caspase-1 was a kind gift from Dr. Gabriel Núñez (University of Michigan Medical School). Ultra-pure LPS (tlrl-pb5lps), Nano-SiO2 (tlrl-sio), poly(dA:dT)/lyovec (tlrl-patc) were purchased from InvivoGen. Nigericin (481990) and EDTA-free protease inhibitor cocktail (11873580001) were purchased from Sigma. *Salmonella* strain SL1344 was originally from Dr. Denise Monack (Stanford University). The constructs for NLRP3-Flag, NLRP3-SFP, and NEK7-HA have been previously described (17).

### Cell culture

iBMDMs from WT C57BL/6 or *Nlrp3*^*-/-*^ mice were maintained in IMDM (Thermo Fisher, 12440053) supplemented with 10% (vol/vol) FBS, L-glutamine, sodium pyruvate, and antibiotics (penicillin/streptomycin). For experiments, iBMDMs were seeded overnight into plates in IMDM with 1% (vol/vol) FBS. HEK 293T cells were cultured in DMEM (Thermo Fisher, 11960044) supplemented with 10% (vol/vol) FBS, L-glutamine, sodium pyruvate, and antibiotics (penicillin/streptomycin). Cultured cells were tested to be free of mycoplasma contamination.

### Generation of mouse Nlrp3^C721Stop/C721Stop^ knock-in macrophages

Alt-R® CRISPR-Cas9 crRNA for mouse *Nlrp3* (targeting sequence: CACGGCAGAAGCTAGAAGTGAGG), tracrRNA (1072533), HDR donor oligos (Sense: TTCTGGCCTCCTCCTTTGCCATTCTAGACTGGTGAACTGCTGACTGACTTCTAGCTTCTGCCGTGGTCTCTTCTCAAGTCTAAGCACC; Antisense: GGTGCTTAGACTTGAGAAGAGACCACGGCAGAAGCTAGAAGTCAGTCAGCAGTTCACCAGTCTAGAATGGCAAAGGAGGAGGCCAGAA), Cas9 nuclease (1081059) and HDR Enhancer (1081072) were purchased from IDT and the RNP complexes were prepared according to the manufacturer’s protocol. The RNP complexes were delivered into iBMDMs by electroporation with nucleofector kit V (Lonza, VCA-1003). Cell clones harboring *Nlrp3*^*C721Stop*^ homologous alleles were identified by sequencing.

### Inflammasome activation assay

iBMDMs were seeded overnight into 12-well plates at a density of 5× 10^5^ per well in IMDM containing 1% (vol/vol) FBS. Cells were primed with 200 ng ml^-1^ ultrapure LPS for 4 h in serum-free IMDM. After priming, cells were then stimulated with ATP (5 mM, 1 h), nigericin (5 μM, 1 h), Nano-SiO_2_ (200 μg ml^-1^, 4 h), Imiquimod (20 μg ml^-1^, 4 h), poly(dA:dT) (2 μg ml^-1^, 4 h), or *Salmonella* [multiplicity of infection (m.o.i.) = 10, 2h]. After stimulation, culture supernatants were collected and cells were directly lysed with 150 μl of 2× laemmli buffer (Bio-rad, 1610737). Proteins in culture supernatants and cell lysates were separated by SDS-PAGE and transferred onto PVDF membranes by using the Trans-Blot Turbo system (Bio-rad). Membranes were immunoblotted with anti-caspase-1, anti-GSDMD, or other indicated antibodies. When specified, equal amounts of cell lysates and supernatants were combined for immunoblotting. The immunoblotting images were developed with hyblot ES^®^ high sensitivity autoradiography films (Denville Scientific, E3218) or captured digitally by using ChemiDoc Imaging System (Bio-rad). The release of cytokines IL-1β and TNF-α into supernatants was determined with ELISA kits (R&D Systems, DY401-05 and DY410-05) according to the manufacturer’s instructions.

### Reconstitution of the full-length NLRP3 and its truncation mutants in iBMDMs

SFP-tagged or HA-tagged full-length mouse NLRP3 or its truncation mutant [NLRP3(1-686), NLRP3(1-720), or NLRP3(730-1033)] were cloned into lentiviral plasmid pHIV-EGFP (Addgene, 21373). Each pHIV-EGFP construct was co-transfected with package plasmids pCMV-VSV-G (addgene, 8454) and pCMV-dR8.2 dvpr (Addgene, 8455) into HEK 293T cells for 96 h. lentiviruses in the culture supernatants were concentrated by using lenti-X concentrator (Takara bio, 631231). iBMDMs were seeded into 6-well plates overnight at a density of 1×10^5^ per well. Cells were then transduced with lentiviruses in the presence of 8 μg ml^-1^ polybrene (Sigma, H9268) for 24 h before the replacement of the fresh culture medium. After 3-4 days, transduced cells were sorted by flow cytometry using GFP as a marker. The expression of reconstituted proteins was determined by immunoblotting with an anti-HA, anti-NLRP3, or anti-Flag antibody.

### Transfection, Immunoprecipitation, and Pull-down assay

HEK 293T cells were seeded into 6-well plates overnight at a density of 6.25 × 10^5^ cells per well. Plasmids expressing HA-tagged NEK7, Flag, or SFP-tagged wild-type NLRP3, or its truncation mutant were single- or co-transfected into HEK 293T cells by Lipofectamine LTX (ThermoFisher, 15338100) for 16 h. After transfection, cells were washed with ice-cold PBS twice and lysed in ice-cold lysis buffer [50 mM Tris-HCl (pH 7.4), 2 mM EDTA, 150 mM NaCl, 0.5% (vol/vol) Nonidet P-40, 1× EDTA-free protease inhibitor cocktail) at 4°C for 10 min. Cell debris was removed by centrifugation at 12,000 ×g for 10 min at 4°C. Cell lysates were precleared by incubation with protein G agarose beads for 1 h at 4°C. Precleared lysates were then incubated with an anti-Flag (1:200) antibody at 4°C overnight. The proteins bound by the antibody were pulled down with protein G beads and subjected to immunoblotting analysis.

### ASC speck staining and imaging

iBMDMs were seeded overnight into an 8-well permanox chamber slide at a density of 1 × 10^5^ cells per well. Cells were primed with 200 ng ml^-1^ LPS for 4 h in serum-free IMDM and then stimulated with 5 mM ATP (30 min), 5 μM nigericin (1 h), or *Salmonella* (m.o.i=10, 4 h). After stimulation, cells were washed once with PBS and then fixed in 4% paraformaldehyde for 20 min at room temperature. Fixed cells were permeabilized with 0.2% Triton X-100 for 5 min, and blocked with PBS buffer containing 5% BSA for 1 h. Cells were incubated overnight with an anti-ASC antibody (1:200) in the blocking buffer. After washing three times with PBS, cells were incubated for 1 h with Alexa-Fluor-488 conjugated anti-rabbit secondary antibody (1:5,000) in the blocking buffer. Cells were washed with PBS three times and mounted with Prolong Diamond Antifade Mountant with DAPI (Thermo Fisher, P36965). Cell images were taken using a Nikon E-800 microscope system and processed with ImageJ.

### ASC oligomerization assay

iBMDMs were seeded overnight in 6-well plates at a density of 1 × 10^6^ cells per well. Cells were primed with 200 ng ml^-1^ LPS for 4 h in serum-free IMDM and then stimulated with 5 mM ATP (30 min), 5 μM nigericin (1 h), or *Salmonella* (m.o.i=10, 4 h). After removing the medium, cells were directly lysed in wells for 15 min at 4 °C with 300 μL of ice-cold PBS buffer supplemented with 0.5% Triton X-100, 0.5 mM PMSF, and 1× EDTA-free protease inhibitor cocktail. Cell lysates were collected by scrapping from each well and transferred to Eppendorf tubes. Lysates were separated into supernatants (TritonX-100-soluble fraction) and pellet (TritonX-100-insoluble fraction) by centrifugation at 6,000 × g for 15 min at 4 °C. The TritonX-100-insoluble fractions were washed with PBS twice and cross-linked for 30 min at room temperature with 2 mM bis[sulfosuccinimidy] suberate (BS^3^) (Thermo Fisher, 21580). The cross-linked pellets were spun down at 6,000 × g for 15 min and dissolved in an SDS sample buffer for subsequent immunoblotting analysis.

### Blue native PAGE and 2D PAGE

Blue native gel electrophoresis was performed as previously described (41). Briefly, iBMDMs were seeded overnight into 6-well plates at a density of 1 × 10^6^ cells per well and stimulated as indicated. After stimulation, cells were washed once with cold PBS and then lysed in ice-cold native lysis buffer (20 mM Bis-tris, 500 mM ε-aminocaproic acid, 20 mM NaCl, 10% (w/v) glycerol, 0.5% digitonin, 0.5 mM Na_3_VO_4_, 1 mM PMSF, 0.5 mM NaF, 1× EDTA-free Roche protease inhibitor cocktail, pH 7.0) for 15 min on ice. Cell lysates were clarified by centrifugation at 20,000g for 30 min at 4°C; Proteins were separated in 4–12% blue native PAGE and then analyzed by western blot. For the two-dimensional (2D) PAGE, the natively resolved gel slice was loaded into the well of 4–12% SDS–PAGE gel after being soaked in 1× Laemmli buffer as previously described (41).

### Statistical analysis

Data are represented as mean ± SEM. Statistical analysis was performed using unpaired two-tailed Student’s t-test or one-way ANOVA with GraphPad 5.0. A *p* value less than 0.05 was considered statistically significant.

## Data availability

All data generated for this study are included in this article.

## Acknowledgments

We thank all the members of the He laboratory for their comments and suggestions. We thank Dr. Gabriel Núñez (the University of Michigan) for providing an anti-caspase-1 antibody, iBMDMs, and other important reagents. We would also like to thank Dr. Jessica Back and Eric Van Buren at the Microscopy, Imaging, and Cytometry Resources Core of Wayne State University for cell sorting.

## Author contributions

Y. D. and Y. H., conceptualization; Y. D. and J.W., methodology; Y. D., J. W., J.C., N.K., investigation; Y. D., J.W., and Y.H. formal analysis; Y.D. and Y.H. writing–original draft. Y.H. funding acquisition; Y.H. supervision.

## Funding and additional information

The work from our laboratory is supported by the US National Institutes of Health (AI148544 to Y. H.) and the Wayne State Startup funds (to Y. H.). The Microscopy, Imaging, and Cytometry Resources Core are supported in part by NIH Center grant P30 CA22453 to the Karmanos Cancer Institute and R50 CA251068-01 to Kamiar Moin, Wayne State University. The content is solely the responsibility of the authors and does not necessarily represent the official views of the National Institutes of Health.

## Abbreviations

iBMDMs: immortalized mouse bone marrow-derived macrophages
NLRP3: NOD-like receptor family pyrin domain-containing 3
LRR: leucine-rich repeats
NATCH: NAIP, CIITA, HET-E, and TP1domain
PYD: pyrin domain
ASC: apoptosis-associated speck-like protein containing a caspase-activation and recruitment domain
GSDMD: gasdermin D
LPS: lipopolysaccharide
m.o.i.: multiplicity of infection
SFP: S-tag, FLAG, and streptavidin-binding tag.

